# Bacterial outer membrane vesicles provide an alternative pathway for trafficking of type III secreted effectors into epithelial cells

**DOI:** 10.1101/415794

**Authors:** Natalie Sirisaengtaksin, Eloise J. O’Donoghue, Sara Jabbari, Andrew J. Roe, Anne Marie Krachler

## Abstract

Outer membrane vesicles (OMVs) are proteo-liposomes universally shed by Gram-negative bacteria. Their secretion is significantly enhanced by the transition into the intra-host milieu and OMVs have been shown to play critical roles during pathogenesis. Enterohemorrhagic *Escherichia coli* O157 (EHEC), causes diarrheal disease in humans, and soluble toxins including Shiga-like toxins that contribute to disease severity and clinical complications including hemolytic uremic syndrome, have been shown to be OMV associated. In addition to Shiga-like toxins, EHEC produces a type III secretion system (T3SS), and T3SS effectors are associated with colonization and disease severity *in vivo*. Here, we show that type III secreted substrates including translocators and effectors are incorporated into OMVs independent of type III secretion activity. EHEC strains with non-functional type III secretion systems shed more OMVs and vesicles enter host cells with accelerated kinetics compared to vesicles shed from wild type EHEC. The T3SS effector translocated intimin receptor (Tir) is trafficked from OMVs into host cells and localizes to the membrane. However, its clustering on the host membrane and co-localization with bacterial pedestals is intimin-dependent. We further show that OMV-delivered Tir can cross-complement an effector-deficient EHEC strain, demonstrating that OMV-associated effectors reach the host cell in a biologically intact form. Finally, we observe that the non-LEE encoded E3 ubiquitin ligase effector NleL is also trafficked to host cells via OMVs, where it ubiquitinylates its target kinase JNK. Together, these data demonstrate that trafficking of OMV-associated effectors is a novel and T3SS-independent pathway for the delivery of active effectors to host cells.

## INTRODUCTION

Enterohemorrhagic *Escherichia coli* (EHEC) are a leading cause of food-borne diarrheal disease world-wide. In some cases, gastrointestinal symptoms can be complicated by the development of hemolytic uremic syndrome (HUS), which is linked with increased morbidity and mortality (1). Secreted toxins, chiefly Shiga-like toxins, lead to the development of HUS, and the accessory toxins cytolethal distending toxin V (CdtV) and hemolysin (Hly) are thought to contribute to HUS pathology (2, 3). However, many more virulence factors are associated with primary colonization of the gastrointestinal tract, in particular the locus of enterocyte effacement (LEE), which encodes for a type III secretion system (T3SS) and associated effectors (4, 5). The T3SS is a needle-like conduit that translocates effector proteins into the host cell where they manipulate host cellular signaling machinery, induce cytoskeletal rearrangements and modulate immunity to facilitate infection. The T3SS needle is formed by the structural protein EspA (6), and secretion activity is driven by the ATPase EscN (7). EscN directly interacts with T3SS chaperones as well as secreted effectors (8). The T3SS initially translocates structural proteins, followed by the translocon components EspD and EspB which perforate the host cell membrane (9, 10), and in response to environmental signals, eventually switches to secrete effectors targeting host proteins (11, 12).

The translocon components EspB and EspD form a complex (13) that embeds into and forms a pore in the host membrane bilayer (14, 15). EspB is both part of the translocon as well as an effector. It has been shown to also be targeted to the host cell cytoplasm (10), where it interacts with catenin and myosin to cause actin reorganization and contribute to microvillus effacement and inhibition of phagocytosis (16, 17). Other translocated effectors include the translocated intimin receptor (Tir), which inserts into the host membrane and contains both intracellular and extracellular domains. The protein is clustered in the membrane via interactions with the adhesin intimin (18), and Tir clustering leads to downstream signaling events that ultimately result in actin reorganization and pedestal formation (18, 19). Tir is critical for colonization and disease pathogenesis in the infant rabbit model (20).

In addition to type III secretion systems, which are pathogen-specific, Gram-negative bacteria also use a range of generalized secretion systems to transport cargo. Outer membrane vesicles (OMVs) are thought to be an alternative secretion system that can facilitate both inter-bacterial (21, 22) as well as bacteria-to-host cargo transfer (23). OMVs are proteo-liposomes of 10-200 nm diameter formed by budding of the outer membrane, and can contain membrane-associated and soluble proteins, nucleic acids and small molecules. OMV production is a constitutive process, but is also a protective response and increases in the presence of environmental stresses and during infection (24-27). Often, this increase in OMV production is co-regulated with other virulence mechanisms (24, 28).

EHEC-derived OMVs play important roles during infection: they constitute a decoy protecting bacteria against antimicrobial peptides (24) and have been shown to enter host cells (29). EHEC OMVs traffic a wide range of cargoes into host cells, including lipopolysaccharide (30, 31), Shiga-like toxins, CdtV and Hly (29, 32), which have been shown to reach the target cells in a biologically active form and contribute to pathogenesis (29, 31). However, localization and trafficking of T3SS components by OMVs has not been studied. Here, we set out to investigate whether T3SS components could be trafficked by OMVs, and whether this process may act as an alternative pathway to facilitate effector delivery to host cells.

## RESULTS

### Type III secreted proteins localize to outer membrane vesicles in the presence and absence of functional type III secretion

First, we set out to determine whether EHEC OMVs contain proteins that are usually secreted through the type III secretion machinery. Type III secretion is a hierarchical process, where needle components are secreted as early substrates, followed by secretion of the translocon protein EspD and EspB, and finally the effectors. We measured the amount of the translocator EspB as well as the T3SS effector translocated intimin receptor (Tir) in total cell lysates, culture supernatants, and purified OMVs from EHEC wild type strain NCTC 12900 and the isogenic secretion deficient mutant Δ*escN*, which lacks the translocation ATPase. We also probed fractions of EHEC wild type strain TUV 93-0 and the isogenic mutant ΔOI-148A. ΔOI-148A contains a deletion of the first half of the LEE pathogenicity island, which removes the important regulator Ler required for expression of the T3SS (33). As previously shown, all strains except the ΔOI-148A mutant produced EspB and Tir (Figure 1). As expected, secretion was only detected in both wild type strains, but not in the Δ*escN* and ΔOI-148A mutants. Purified OMVs contained both translocon components EspB and EspD, as well as the effector Tir, and localization of the T3SS components to OMVs required the presence of the T3SS locus, but did not depend on a functional T3SS, since OMVs collected from the Δ*escN* mutant also contained EspB, EspD and Tir (Figure 1).

**Figure 1.**
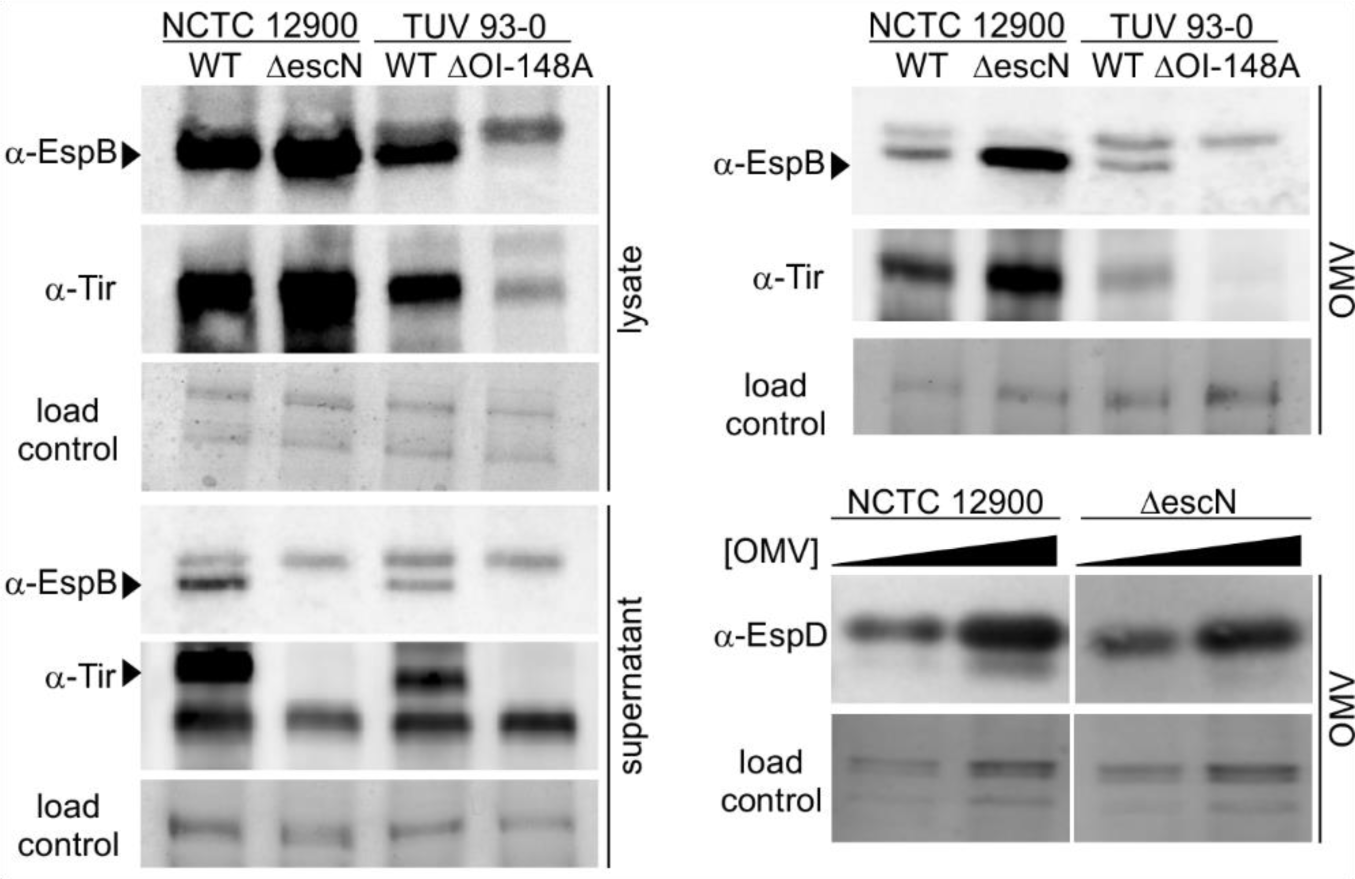
Type III secreted proteins localize to outer membrane vesicles in the presence and absence of functional type III secretion. Total cell lysates, supernatants, and OMVs were harvested from NCTC 12900 wt and an isogenic Δ*escN* deletion strain, or a TUV 93-0 wt and an isogenic ΔOI-148A deletion strain. All fractions were normalized using OD_600_ readings at harvest. For EspD blots, OMV concentrations were measured, and adjusted to concentrations of 10^11^ (left lanes) and 2*10^11^ OMVs/ml (right lanes) for both strains. All fractions were separated by SDS-PAGE and either probed by Western Blotting using α-EspB, α-Tir, or α-EspD antibodies to visualize the translocon components EspB and EspD and the effector translocated intimin receptor (Tir), or stained with Coomassie Blue to provide loading controls.

### A type III secretion deficient EHEC strain produces excess outer membrane vesicles

To determine the effect of T3SS activity on OMV production, EHEC wild type and a T3SS secretion deficient strain, Δ*escN* were compared. The Δ*escN* strain produces T3SS structural components and effectors, but is unable to secrete them since the associated T3SS ATPase is lacking (34). Both strains were grown in LB for 18 hours, culture densities were normalized, and OMVs were harvested and quantified using Nanosight particle tracking analysis. The Δ*escN* mutant produced significantly more OMVs per cell than the NCTC 12900 wild type strain (Figure 2A). The mean diameter of vesicles produced by the isogenic Δ*escN* strain was slightly larger than of OMVs derived from the corresponding wild type strain (Figure 2B). These data show that in the absence of a functional type III secretion system, EHEC hypervesiculates and produces OMVs of aberrant morphology.

**Figure 2.**
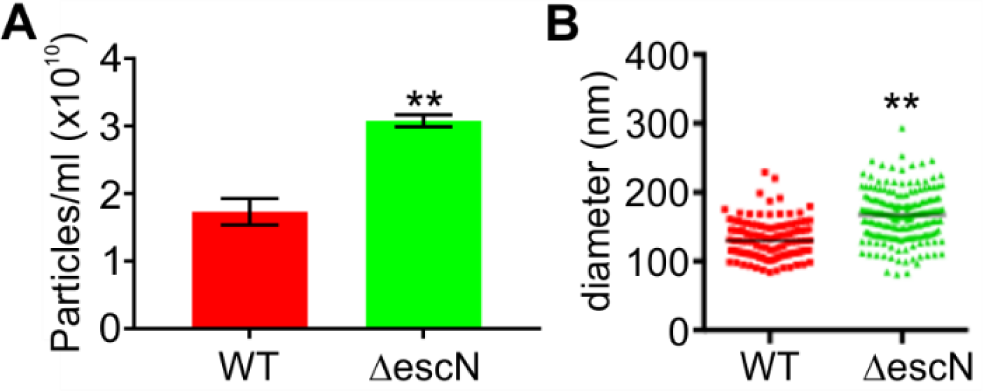
A type III secretion deficient EHEC strain produces excess outer membrane vesicles. OMVs were isolated from an equal biomass of NCTC 12900 wt (red) or Δ*escN* (green) cultures grown in LB for 18 hrs at 37 °C. OMV concentrations **(A)** and OMV size distributions **(B)** were measured using a Nanosight LM10 particle tracking system. A minimum of 100 tracks per sample were determined for three individual preparations per strain. Statistical significance was determined using unpaired t-tests and is depicted with asterisks. (**) indicates a p-value ≤0.01.

### Outer membrane vesicle uptake by host cells is accelerated in the absence of a functional type III secretion system

To determine if loss of a functional type III secretion system would affect the kinetics of OMV entry into host cells, we used a CCF2-AM based reporter assay as previously described (30). This assay uses a vesicle-targeted beta-lactamase reporter (ClyA-Bla) to measure the fast entry kinetics of OMVs into the host cell cytoplasm in real time. Epithelial host cells were pre-loaded with the fluorescent beta-lactamase substrate CCF2-AM, which upon enzymatic cleavage shifts from green to blue fluorescence emission (30, 35). The ClyA-Bla reporter was expressed in either EHEC NCTC 12900 wild type cells or the isogenic Δ*escN* mutant. OMVs were harvested and adjusted to a multiplicity of infection (MOI) of 1000 OMVs per cell, using concentration data from particle tracking analysis. As previously determined (30), this corresponds to an MOI of approx. 37 bacteria/cell, a physiologically relevant level. OMVs were incubated with dye-loaded epithelial cells and uptake kinetics were followed for three and a half hours (Figure 3A, B). Both wild type and Δ*escN* derived OMVs were rapidly taken up by host cells, but OMVs from Δ*escN* were taken up with higher efficiency (Figure 3C) and significantly faster (Figure 3D) than vesicles derived from the wild type strain. We conclude that in the absence of a functional T3SS, OMV uptake is accelerated. Since we have previously established that the chemical structure of the bacterial lipopolysaccharide plays a significant role in rate and efficiency of OMV uptake (30), we analyzed the LPS structure of NCTC 12900 wild type and Δ*escN* strains. As a control, we also analyzed the LPS of the EHEC strain TUV 93-0, and a corresponding isogenic mutant (ΔOI-148A) that does not express the T3SS (33). While the short and very long O-antigen chains were similar in all four backgrounds, both the Δ*escN* and ΔOI-148A featured slightly more long O-antigen chains than either wild type strain (Figure 3E, arrow). This suggests that in the absence of a functional T3SS, the lipopolysaccharide composition is slightly altered.

**Figure 3.**
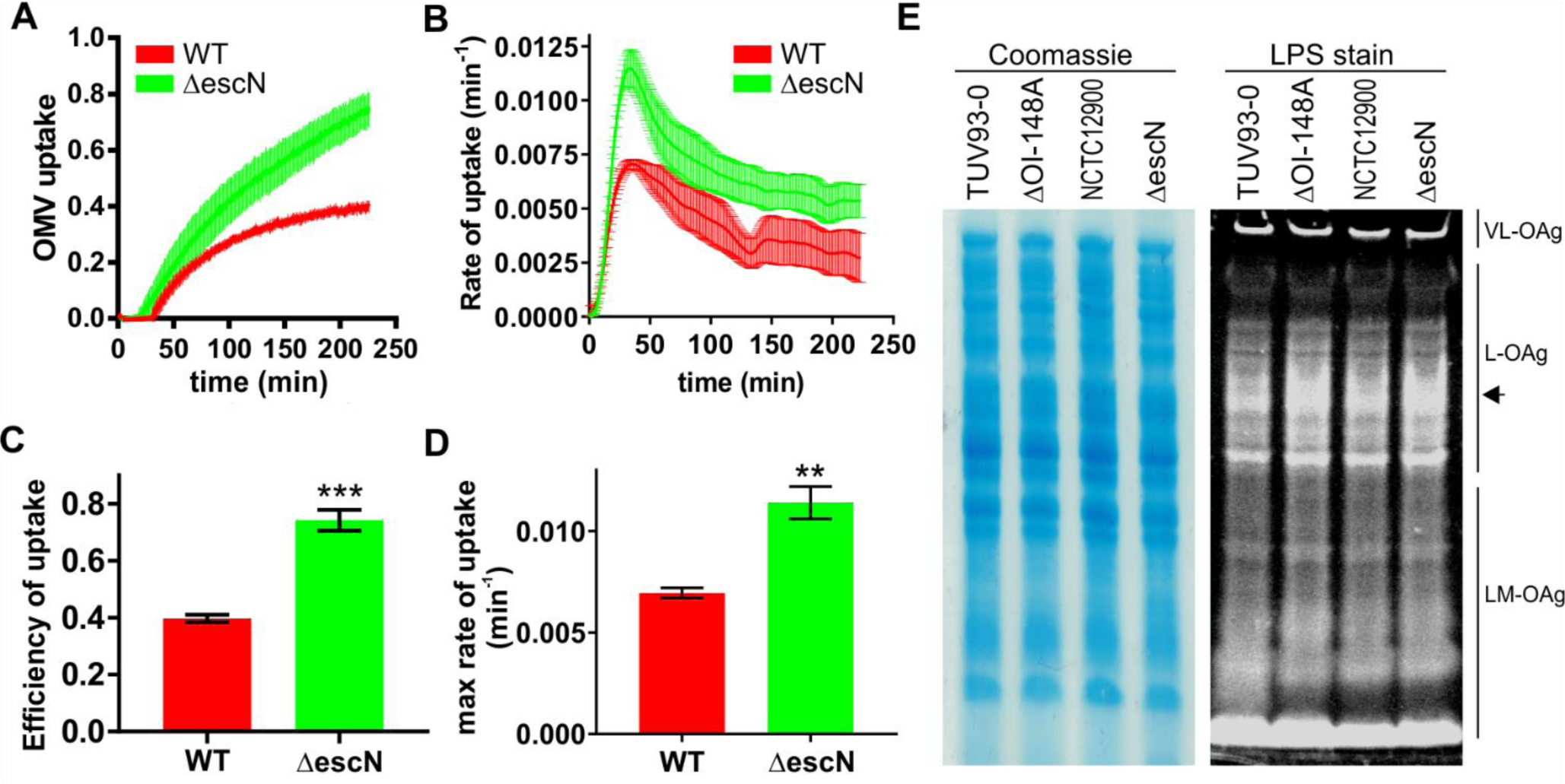
Outer membrane vesicle uptake by host cells is accelerated in the absence of a functional type III secretion system. (A) CCF2-AM loaded Hela epithelial cells were exposed to OMVs from EHEC NCTC 12900 wt (red) or a Δ*escN* deletion strain (green) carrying ClyA-Bla, at an MOI of 1000 for 3.5 hours. Ratios of blue:green fluorescence over time were plotted as means ± sem (n=3). Rates **(B)** were extracted from data in A to visualize speed of OMV uptake over time and are means ± sem (n=3). FRET signal changes after 3.5 hrs (uptake efficiency), **(C)** and maximum rates of uptake **(D)** were determined from data in (A) and (B), respectively. Data shown are means ± sem (n=3). Significance was determined by unpaired t-test, with (**)p≤0.01 and (***) p≤0.001. Cultures of EHEC TUV 93-0 wt or T3SS deficient mutant ΔOI-148A, NCTC 12900 wt or T3SS secretion deficient mutant Δ*escN* were separated by SDS-PAGE and total protein and LPS were visualized by Coomassie Brilliant Blue and Pro-Q Emerald 300 staining, respectively. VL-OAg: very long O-antigen, L-OAg: long O-antigen, LM-OAg: low and medium length O-antigen. Arrow on right marks region of difference between wild type and mutant strains.

### Outer membrane vesicles get delivered to host cells independent of type III secretion activity

Although the use of purified OMVs for infection experiments allowed us to normalize the number of vesicles used, the system is somewhat artificial, in that it does not follow the physiological time course of vesicle formation, release, and diffusion to host cells. To more closely mimic the time line of vesicle-based trafficking during an infection, we set up a transwell experiment, where bacteria were grown in the transwell top compartment to allow OMV production and release, and host cells were cultured in the bottom compartment to permit contact with OMVs following diffusion through the transwell, but exclude access and direct contact with intact bacterial cells (Figure 4A). Uptake of OMVs by host cells was then measured using CCF2-AM loaded epithelial cells and ClyA-Bla expressing EHEC strains. OMV uptake was continuously observed over 15 hours, and although as expected, the uptake kinetics were significantly slower than those observed with isolated OMVs (Figure 4B), the pattern was consistent with results obtained using isolated vesicles: OMVs secreted by the wild type strain were taken up more slowly (Figure 4C) and less efficiently (Figure 4D) than those derived from the Δ*escN* mutant.

**Figure 4.**
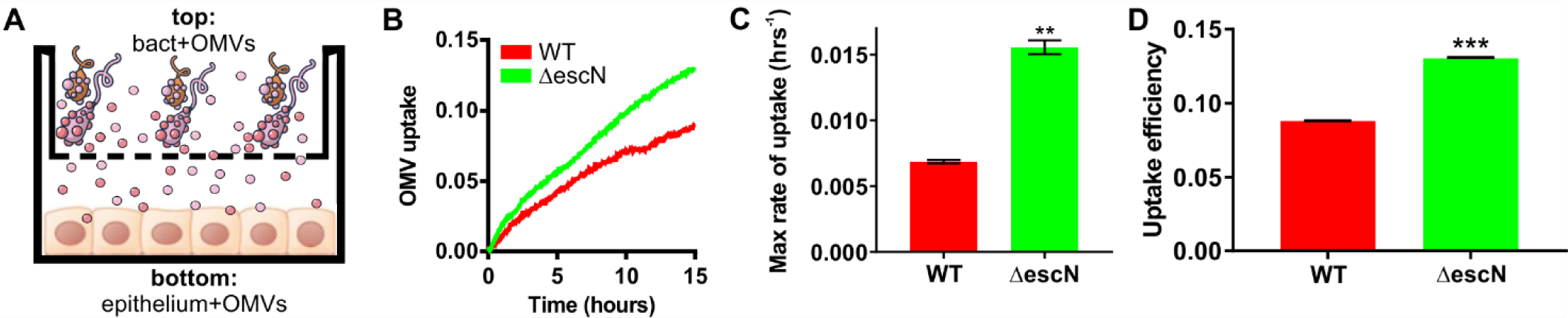
Outer membrane vesicles get delivered to host cells independent of type III secretion activity. **(A)** Experimental setup for OMV release and delivery experiments. Epithelial cells are grown in the bottom compartment of a transwell and loaded with CCF2-AM dye. Bacteria (MOI=10) are added to the top compartment of the transwell, and OMV uptake by host cells in the bottom well is monitored by measuring FRET over time. (B) CCF2-AM loaded Hela cells were exposed to EHEC NCTC 12900 wt (red) or a Δ*escN* deletion strain (green) carrying ClyA-Bla, at an MOI of 10 for 15 hours. Ratios of blue:green fluorescence over time were plotted as means ± stdev (n=3). Maximum rates **(C)** were extracted from data in Fig.4B to visualize speed of OMV uptake and are means ± stdev (n=3). FRET signal changes after 15 hrs **(D)** were determined from data in (B) and plotted to visualize overall efficiency of uptake. Data shown are means ± stdev (n=3). Significance was determined by unpaired t-test, with (**) p≤0.01 and (***) p≤0.001.

### The T3SS effector translocated intimin receptor (Tir) is delivered to host cells via OMVs

Since we established that T3SS effectors associate with OMVs, we set out to test if OMV associated effectors would be trafficked into host cells. Intestinal epithelial (RKO) cells were infected with wild type or secretion deficient (Δ*escN*) bacteria, or wild type and T3SS deficient (ΔOI-148A) derived OMVs for four hours, at an MOI of 30 bacteria/cell or a corresponding MOI of 1000 OMVs/cell. Host cells were then fixed and stained with α-Tir antibody and Hoechst to visualize translocated intimin receptor within host cells, and DNA, respectively (Figure 5).

**Figure 5.**
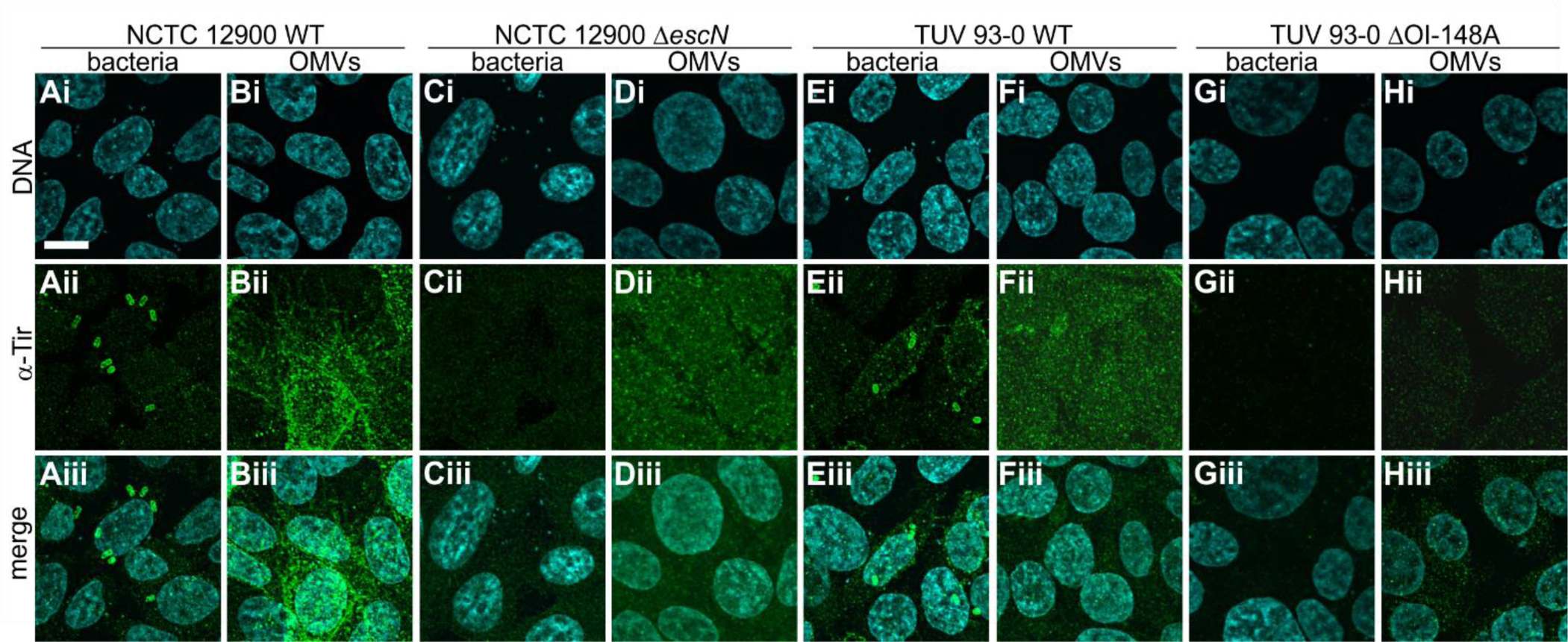
The T3SS effector translocated intimin receptor (Tir) is delivered to host cells via OMVs. Intestinal epithelial (RKO) cells were infected with bacteria at an MOI of 30 (A, C, E, G) or purified OMVs at an MOI of 1000 (B, D, F, H) for 4 hours. Strains used were NCTC 12900 wild type (A, B), NCTC 12900 Δ*escN* (C, D), TUV 93-O wild type (E, F) and TUV 93-O ΔOI-148A (G, H). RKO cells were fixed in PFA, permeabilized with Triton X-100, and stained. Images of Hoechst to visualize DNA (i), α-Tir antibody to visualize translocated intimin receptor (ii) and merged images (iii) are shown. Images are representative of n=3 experiments. Scale bar, 10 μm.

In intestinal epithelial cells infected with wild type bacteria (NCTC 12900, Fig. 5A or TUV 93-0, Fig. 5E), Tir was found throughout the host cell, and was particularly concentrated in foci (pedestals) below attached bacteria. In contrast, Tir was absent from cells infected with bacteria deficient in T3SS secretion (Δ*escN*, Fig. 5C) or deficient in T3SS expression (ΔOI-148A, Fig. 5G). When cells were infected with OMVs derived from both wild type strains as well as the secretion deficient mutant Δ*escN*, Tir was present in host cells. However, its localization was not focal but diffuse all over the plasma membrane (Fig. 5B, D, F). Tir was absent following incubation with OMVs derived from the ΔOI-148A strain which does not express the T3SS (Fig. 5H). We also tried to immunostain cells using an α-EspB antibody, but the signal was too weak to draw conclusions, even after prolonged incubation. Together, our data suggests that effectors usually secreted via the EHEC T3SS can also be delivered to intestinal cells via outer membrane vesicles.

### OMV-delivered translocated intimin receptor reaches the host cell in a biologically active form

Since we observed that the T3SS effector Tir could be trafficked into intestinal cells via OMVs, we next asked whether Tir would reach the host cell in a biologically active form. To test the biological activity of OMV-derived Tir, we infected intestinal epithelial cells with a Tir-deficient EHEC strain, which attaches to host cells but fails to form pedestals (Fig. 6A). We then asked whether Tir-containing OMVs could cross-complement and reconstitute pedestal formation for the Tir-deficient strain. Tir was absent from cells infected with EHEC *Δtir* (Fig. 6A), but present when infections were carried out in the presence of OMVs derived from wild type or Δ*escN* strains (Fig. 6B, C). The presence of bacteria caused the redistribution of OMV-derived Tir, which formed foci under the bacterial cells (Fig. 6B, C). No Tir or pedestals were observed for uninfected cells (Fig. 6D). Overall, bacterial attachment was enhanced and actin pedestal formation restored in the presence of OMV-derived Tir (Fig. 6E, F).

**Figure 6.**
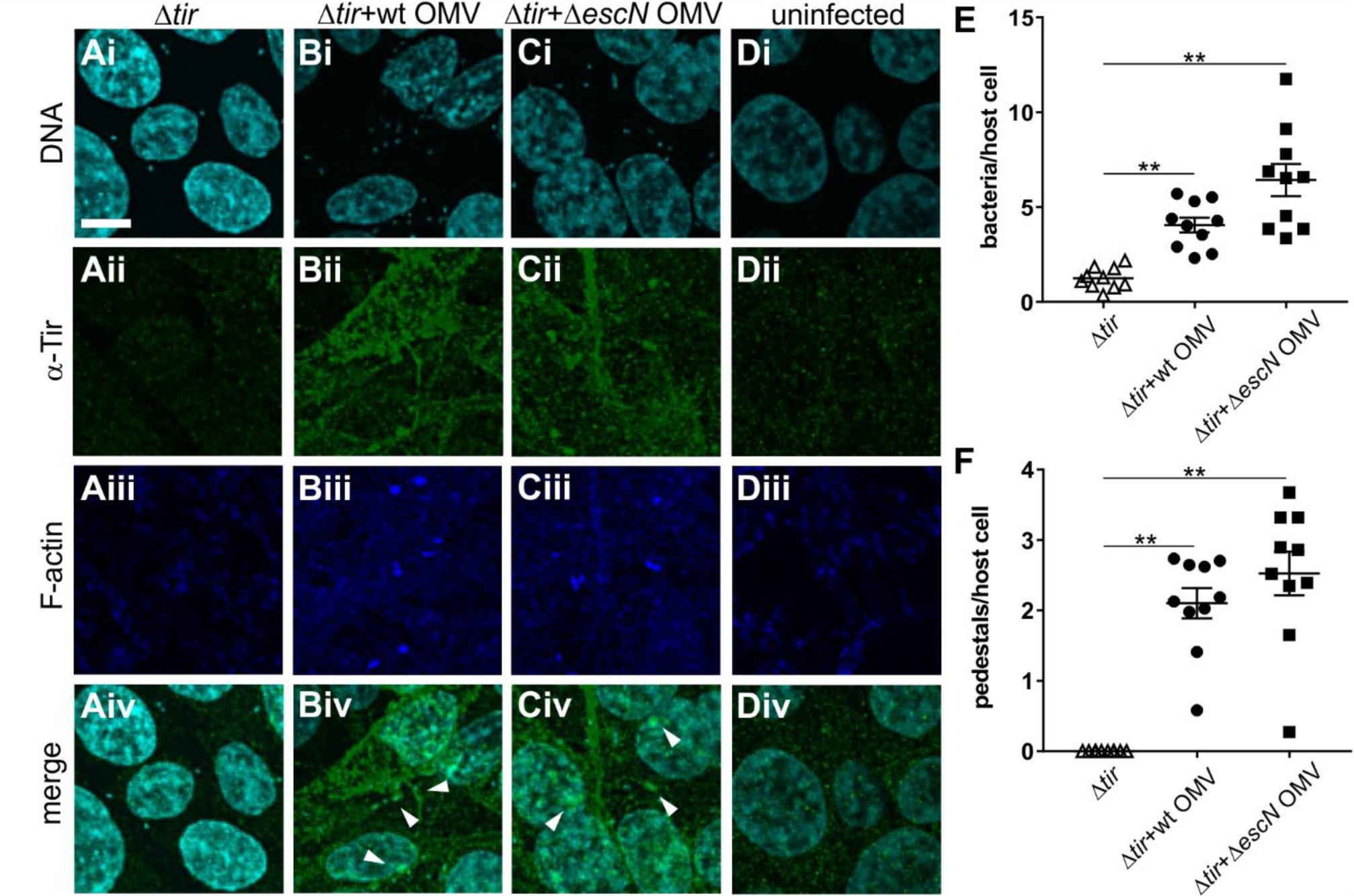
OMV delivered Tir can restore pedestal formation by Tir deficient EHEC. Intestinal epithelial (RKO) cells were infected for 4 hours with bacteria (EHEC Δtir) at an MOI of 30 either alone **(A)**, or in the presence of OMVs at an MOI of 1000 derived from NCTC 12900 wild type **(B)** or Δ*escN* mutant **(C)**. As a control, RKO cells were left uninfected **(D)**. RKO cells were fixed in PFA, permeabilized with Triton X-100, and stained. Images of Hoechst to visualize DNA (i), α-Tir antibody to visualize translocated intimin receptor (ii), phalloidin to stain F-actin (iii) and merged images (iv) are shown. Images are representative of n=3 experiments. Scale bar, 10 μm. **(E)** Bacteria per host cell and **(F)** pedestals per host cell were quantified from imaging experiments. Individual counts, means and s.e.m. from 10 slides (a total of ≥500 RKO cells) across three independent experiments are shown. Significance was analyzed using one-way ANOVA and Brown-Forsythe test. (**) indicates a corrected p-value ≤ 0.01.

### EHEC-derived OMVs traffic non-LEE encoded effectors into host cells

In addition to LEE-encoded structural components and effectors, the EHEC T3SS also secretes a number of non-LEE encoded effectors that are scattered throughout the EHEC chromosome. One such effector is the non-LEE encoded effector ligase (NleL), a E3 ubiquitin ligase that ubiquitinylates and inactivates the host c-Jun N-terminal kinases (JNKs) (36). To test whether non-LEE effectors could be trafficked within EHEC-derived OMVs, we infected host cells with different concentrations of OMVs or intact NCTC 12900 cells at an MOI of 30, harvested cell lysates and probed Western Blots with anti-JNK antibody. While with untreated host cells, three bands characteristic of different JNK isoforms were observed, cells infected with EHEC feature an additional band shifted by approx. 10 kDa, corresponding to the molecular weight of one ubiquitin subunit (8.5 kDa). The addition of EHEC-derived OMVs to host cells similarly led to the concentration-dependent appearance of ubiquitinylated JNK (Fig. 7). These data suggest that NleL, a non-LEE encoded T3SS effector, is also trafficked to host cells in a biologically active state via outer membrane vesicles.

**Figure 7.**
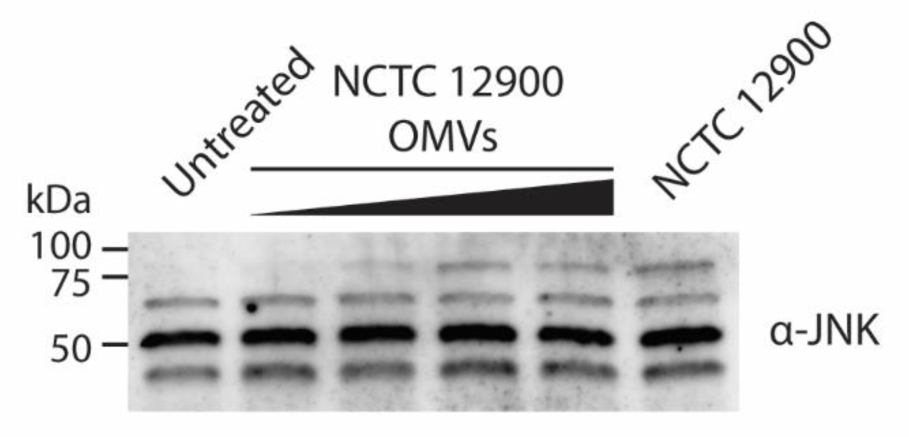
RKO cells were left untreated, or infected with increasing amounts of NCTC 12900-derived OMVs, or NCTC 12900 at an MOI of 30, for 4 hours. Cells were harvested, lysed, lysates separated byu SDS-PAGE, Western Blotted, and probed using an anti-JNK antibody.

## DISCUSSION

Outer membrane vesicles are abundantly produced by all Gram-negative bacteria, and OMV production has been shown to enhance the pathogenic potential of bacteria, including EHEC. OMVs isolated from EHEC O157 have been shown to contain a range of virulence factors, including Shiga-like toxins and hemolysin, which are traditionally thought of as secreted toxins (37, 38). EHEC OMVs are sufficient to cause hemolytic uremic syndrome-like symptoms (39) and are a key activator of pro-inflammatory responses to bacterial infections *in vivo* (31). OMVs additionally play important roles during infection, including as a decoy against host-secreted antimicrobial peptides (24). OMVs are constantly produced, but the transition to the intra-host milieu and exposure to cues such as colonic medium and mucin significantly increases vesiculation. OMV production is highest in host-isolated EHEC and decreases during lab cultivation (40). Similarly, antibiotic treatment increases OMV production and the amount of OMV-associated Shiga-like toxins (40). These quantitative differences between OMV production in laboratory culture and in vivo are likely the reason why OMV mediated trafficking of T3SS effectors has thus far eluded detection in conventional *in vitro* culture assays.

In addition to Shiga-toxins, type III secreted effector proteins are critical factors promoting EHEC colonization and pathogenesis *in vivo* (4). T3SS effectors modulate key cellular signaling processes, leading to cytoskeletal rearrangements and modulation of host immune responses. In this study, we set out to investigate whether type III secreted proteins are also secreted via outer membrane vesicles, and whether they would be biologically active within the host.

OMV biogenesis is thought to require local instabilities in the bacterial cell envelope – either the local loss of linkage between outer membrane and peptidoglycan layer, or local changes in the outer membrane organization, such as changes in phospholipid content or curvature (41). Here, we found that failure to assemble a functional type III secretion system also leads to increased vesiculation (Fig. 2). The ATPase deficient Δ*escN* mutant assembles the type III secretion system basal body, but is incapable of producing a needle and secreting effectors. The T3SS structure traverses both membranes as well as the peptidoglycan layer, and T3SS assembly is known to be accompanied by local remodeling of the peptidoglycan layer (42, 43). It is conceivable that the initiation of peptidoglycan remodeling by T3SS associated lytic transglycosylases without concurrent full assembly of the T3SS leads to local changes in envelope structure that result in increased vesiculation. The mechanistic details of this process have yet to be elucidated.

We report here that EHEC OMVs contain T3SS associated factors (Fig. 1), including the translocon components EspB and EspD, as well as the effector translocated intimin receptor (Tir) – factors which are ordinarily moved from the bacterial cytoplasm to the host cell. Although current models of OMV biogenesis depict vesiculation as a process of outer membrane blebbing, it is well known that OMVs contain cytoplasmic factors as well as periplasmic and outer membrane components. EHEC-derived OMVs have been shown to contain bacterial DNA, including genomic DNA encoding for virulence factors (37). Although the finding that T3SS associated substrates are contained in OMVs is novel, substrates for other secretion systems have previously been found associated with OMVs. For example, OMVs of the Gram-negative oral pathogen *Tannerella forsythia* are enriched with substrates of the type IX secretion system (44). Interestingly, Ernst et al. recently identified a chaperone-independent secretion pathway for T3SS effectors in the intestinal pathogen *Shigella*, although the involved pathway was not subjected to detailed analysis (45). It would be interesting to see whether the identified pathway is consistent with OMV-dependent effector secretion.

We further found here that the uptake of EHEC OMVs derived from a secretion deficient *ΔescN* T3SS mutant was accelerated compared to OMVs from the wild type strain (Fig. 3). We have previously described that the preferred uptake pathway, and thus entry kinetics of OMVs are shaped by the strain’s lipopolysaccharide composition (30). Although we identified minor changes in the long O-antigen of T3SS deficient strains compared to wild type isolates (Fig. 3) which would be consistent with the accelerated uptake kinetics, it is unclear whether those changes are sufficient to explain the accelerated entry kinetics we observed here. It has been described previously that LPS composition and T3SS activity can impact each other (46, 47), and that O-antigen length is inversely correlated with the amount of T3SS expression (48). Additionally, it is possible that the T3SS needle structure facilitates targeting to the clathrin-mediated endocytic route, which mediates slower trafficking to the cytoplasm, and that the absence of the needle structure from both mutants drives preferential uptake via raft-mediated endocytosis, which enables faster and more efficient trafficking of OMV contained substrates to the cytoplasm, as we have described previously (30).

We observed that OMV-associated T3SS effectors reach the host cell cytoplasm, and retain their endogenous biological activity in the context of an infection (Fig. 5 and 6). The delivery of biologically active virulence factors we observe here is in agreement with other studies that found OMV-delivered Shiga toxin, hemolysin and cytolethal distending toxin V all retain biological activity within host cells, despite being subject to endosomal trafficking (29). We found that OMV-delivered Tir preferentially localizes to the host cell membrane, both when translocated through the T3SS and when delivered via OMVs (Fig. 5).However, the characteristic focal pattern we observe upon whole cell infection (Fig. 5A and E) is absent when the effector is delivered via OMVs, where we instead observe diffuse localization on the host plasma membrane (Fig. 5B, D, F). However, co-localization with bacteria and with actin pedestals is restored for vesicle-trafficked Tir, once bacteria attach to the host cell (Fig. 6). This indicates that the effector remains biologically active, but its correct localization is dependent on the presence of attached bacteria, and is most likely induced by the presence of its cognate ligand, intimin. This idea is consistent with previous work on EPEC Tir and intimin, where the formation of ring-like supramolecular structures were induced in Tir-primed host cells by the addition of intimin containing proteoliposomes (49). Similarly, we demonstrated that NleL, a non-LEE encoded effector, is transported to host cells and retains its enzymatic activity as a E3 ubiquitin ligase. As such, it facilitates colonization by modulating actin dynamics (36). It may have secondary effects, suppressing pro-inflammatory and/or pro-apoptotic signaling which are both downstream of JNK.

Together, our data provide evidence for the trafficking of biologically active type III secreted effectors to epithelial cells by outer membrane vesicles, an alternative pathway for the delivery of EHEC effectors to host cells. It is conceivable that this pathway is relevant during infection *in vivo*, where the amount of bacterial vesiculation is significantly higher than observed under laboratory conditions. It has already been demonstrated by several groups that EHEC OMVs significantly contribute to pathogenicity *in vivo* (31, 39). To what extent the OMV-mediated delivery of T3SS substrates to host tissues is important for virulence *in vivo* will be the subject of future studies.

## EXPERIMENTAL PROCEDURES

### Bacterial strains and growth conditions

The strains used in this study were the *E. coli* serotype O157:H7 strain NCTC 12900, a STX negative isolate (NCTC12900), and the isogenic T3SS ATPase deficient Δ*escN* mutant (50, 51), as well as strain TUV 93-0 and isogenic deletions ΔOI-148A and Δ*tir* (33). For studying entry kinetics, bacterial strains were transformed with plasmid pBAD-ClyA-Bla (kan^R^), a kanamycin resistant derivative of the pBAD amp^R^ vector (52) provided by Matthew DeLisa, Cornell University. Strains were grown in LB containing 50 μg/ml kanamycin for selection where necessary, at 37 °C with shaking at 200 rpm. ClyA-Bla expression was induced by adding 0.02% arabinose to the growth medium.

### Isolation of outer membrane vesicles

The detailed procedure for OMV isolation has been published before (30). Briefly, for isolation of OMVs, strains were grown in LB for 18 hrs. Bacterial cells were pelleted by centrifugation at_600_0xg, and supernatants filtered through 0.45 µm filters. Filtrate was plated to ensure sterility. OMVs were pelleted from 25 ml of filtrate by ultracentrifugation at 100,000xg for 2 hours at 4 °C, washed with DMEM, and resuspended in 1ml of sterile DMEM until further use.

### Nanosight analysis of outer membrane vesicles

To measure concentrations, purified vesicles were diluted 1:10^6^ in filtered PBS, and OMV concentrations and diameters were determined using a Nanosight LM10 particle tracking system. A minimum of 100 tracks were determined per sample, with analysis performed on three separate preparations per sample. Original concentrations were calculated and normalized to cell densities. Prior to infection experiments, OMV preparations were diluted in DMEM to normalize them to an MOI of 1000 (corresponding to an MOI of 37 bacteria/cell).

### Western Blotting

EHEC strains were grown in LB for 18 hours at 37 °C. Cells were harvested by centrifugation, normalized to an OD of 10 in 5x SDS-PAGE loading buffer and boiled. 10 µl were analyzed by SDS-PAGE to determine contents of total cell lysates by Western Blotting. Supernatants were concentrated by acetone precipitation, and amounts normalized to cell densities were analyzed by Western Blotting. OMV fractions were concentrated and normalized to give a concentration of 10^10^particles/ml, and 10 µl were analyzed by SDS-PAGE. Gels were transferred to a PVDF membrane, blocked for 1 h at 22 °C in TBST 5% skimmed milk. Membranes were probed with primary antibodies (α-EspD 1:2000), α-Tir (LS-C500576 from LSBio, 1:1000) or α-EspB (ABIN 1098174, 1:1000) in blocking buffer at 4 °C for 16 hrs, and secondary α-mouse HRP (1:5000) for 1 h at 22 °C. Bound antibodies were detected using Pierce ECL Plus Western Blotting Substrate.

### Infection experiments

Hela cells were seeded at 10^5^cells/ml into black walled, clear bottom 96 well plates 24 hours prior to infections. On the day of the experiment, cells were loaded with CCF2AM for one hour at 22 °C as per the manufacturer’s instructions, and then washed and media replaced with supplement-free DMEM. Cells were incubated with OMVs from EHEC to give an MOI of 1000 (corresponding to a cell-based MOI of approx. 37), and blue and green fluorescence were monitored using a PheraStar Omega plate reader, using excitation at 405 nm and simultaneous dual emission at 530 nm and 460 nm, every 2 minutes as previously described (30).

For transwell infection experiments, Hela cells were seeded in 24 well plates in complete DMEM at 10^5^ cells/ml 24 hrs prior to infections. Cultures of EHEC NCTC 12900 and Δ*escN* were grown in LB and induced with 0.02% L-arabinose. 200 µl of cultures were added to the transwell inserts, with 600 µl of DMEM without supplements added to the cells in the bottom compartment. Plates were analyzed using a PheraStar Omega plate reader for 15 hours, monitoring blue and green fluorescence with excitation at 405 nm and simultaneous dual emission at 530 nm and 460 nm. All experiments were performed with a minimum of three technical replicates and three independent repeats. The ratio of blue to green fluorescence intensity detected in the cells at each cycle was calculated for each experiment.

### Rate estimation, efficiency of uptake, and statistical analysis

To estimate the gradients of the data (i.e. rates of uptake), polynomials were fitted to each data set using the cubic spline function *csaps* in Matlab R2017b. Numerical estimates of the gradients of the resulting polynomials were determined using the *gradient* function. To ensure that the gradient estimates were as smooth as possible whilst also retaining the overall shape and trend of the data, a small smoothing parameter was used. Analysis of variance (ANOVA) was used to determine statistical significance, with a Brown Forsythe test to determine equal variance (GraphPad Prism software). A p-value of <0.05 was considered statistically significant. Efficiency of uptake was calculated as the absolute change in blue:green fluorescence intensity ratio between 0 and 3 hours ([Em460/Em530]_t=0hrs_)/[Em460/Em530]_t=3hrs_). Analysis of variance (ANOVA) was used to determine statistical significance, with a Brown Forsythe test to determine equal variance (GraphPad Prism software). A p-value of <0.05 was considered statistically significant.

### Immunofluorescence staining and microscopy

Infections were carried out as described above, with the exception that RKO cells were used and seeded onto glass cover slips. Cells were infected with EHEC or OMVs as described above for four hours, washed with PBS, and fixed with 3.2% paraformaldehyde in PBS at 4 °C for 16 hours. Fixed cells were permeabilized with 1% Triton X-100 in PBS for 10 minutes, and blocked using 1% BSA in TBST for one hour. Cells were then stained with α-Tir (LS-C500576 from LSBio, 1:1000 in blocking buffer) for one hour at room temperature, washed three times with TBST, and stained with α-rabbit secondary antibody conjugated to Alexa-488 (1:500 in blocking buffer) for one hour at room temperature. Cells were washed three times with TBST, stained with Hoechst (1:1000 in PBS) and phalloidin conjugated with Alexa-680 (1:100 in PBS) for 10 minutes and washed three times with PBS and one with water prior to mounting in ProLong Antifade Gold. Slides were imaged after curing in the dark for 16 hours, using an Olympus IX83 with a Fluoview FW3000 confocal system and a 100x oil immersion objective.

## ACKNOWLEDGEMENTS

We thank Matthew DeLisa (Cornell Univ.) for providing ClyA-Bla plasmid constructs. Funding was provided by a UT Systems STAR award and the NIH (R01AI132354) to A.M.K. A.M.K. and S.J. gratefully acknowledge support from the Biotechnology and Biological Sciences Research Council (BBSRC grant BB/M021386/1).

